# Fractality and Entropic Scaling in the Chromosomal Distribution of Conserved Noncoding Elements in the Human Genome

**DOI:** 10.1101/027763

**Authors:** Dimitris Polychronopoulos, Labrini Athanasopoulou, Yannis Almirantis

**Author notes:** Present address: Department of Molecular Sciences, Institute of Clinical Sciences, Faculty of Medicine, Imperial College London and MRC Clinical Sciences Centre, Hammersmith Hospital Campus, Du Cane Road, London W12 0NN, UK.

## Abstract

Conserved, ultraconserved and other classes of constrained non-coding elements (referred as CNEs) represent one of the mysteries of current comparative genomics. These elements are defined using various degrees of sequence similarity between organisms and several thresholds of minimal length and are often marked by extreme conservation that frequently exceeds the one observed for protein-coding sequences. We here explore the distribution of different classes of CNEs in entire chromosomes, in the human genome. We employ two complementary methodologies, the scaling of block entropy and box-counting, with the aim to assess fractal characteristics of different CNE datasets. Both approaches converge to the conclusion that well-developed fractality is characteristic of elements that are either marked by extreme conservation between two or more organisms or are of ancient origin, i.e. conserved between distant organisms across evolution. Given that CNEs are often clustered around genes, especially those that regulate developmental processes, we verify by appropriate gene masking that fractal-like patterns emerge irrespectively of whether elements found in proximity or inside genes are excluded or not. An evolutionary scenario is proposed, involving genomic events, that might account for fractal distribution of CNEs in the human genome as indicated through numerical simulations.

## INTRODUCTION

Long-range correlations were shown to be present in the nucleotide sequence of the non-protein-coding part of eukaryotic genomes, soon after such genomes were sequenced^1–3^. In previous studies, we investigated the distributional features that extend at a large-scale, of genomic elements such as protein coding sequences^4^, transposable elements^5,6^ and conserved noncoding elements^7^, by exploring the size distribution of inter-exon, inter-repeat and inter-CNE distances respectively. In most cases we observed power-law-like size distributions that often span several orders of magnitude.

In information theory, entropy was conceived by Claude Shannon^8^ to be an estimator of the amount of information that is carried in a transmitted message. During the last decades, scale invariance and fractality have been found in time series from signal transmission in electronic engineering, earthquakes, economy, social sciences and many other fields. Very often, such studies have been carried out using the standard box-counting technique and, in several cases of systems characterized by long range correlations, Shannon entropy has also been used in applications including biological sequence analysis^9–11^.

In a previous work^12^ we studied the scaling properties of the block entropy of the distribution of genes in whole chromosomes of eukaryotic genomes through a coarse-graining reduction of the DNA sequence into a symbol sequence. The convention we followed was that zeros “0” in the symbol sequence stood for non-protein-coding nucleotides and ones “1” for nucleotides belonging to Protein Coding Segments (coding exons, denoted as PCSs). Several studies have shown that a linear scaling of the Shannon-like (or block) entropy *H(n)* with the length *n* of the word (called hereafter *n*-word or block of length *n*) in semi-logarithmic plots is a clear indication of long-range order and fractality, as we are going to discuss in the next section^13–16^. We verified this conjecture numerically in the case of finite Cantor-like symbol sequences^12^. Then, we showed that the genomic distribution of protein coding segments often exhibits this particular scaling. In a more recent work^17^, we studied the scaling properties of the block entropy in the chromosomal distribution of Transposable Elements (TEs) and again we found the occurrence of the aforementioned scaling. The observed linearity in semi-logarithmic plots in the two types of genomic components follows different modalities. We have been able to attribute the observed distributional patterns, as expressed by entropic scaling and their differences, to the different evolutionary history of PCSs and TEs by means of a simple model. The model is shown, through computer simulations, to reproduce the observed pattern and includes key evolutionary events characterising both genomic elements highly conserved in the course of evolution (e.g. protein coding segments and CNEs) and genomic elements mostly non-conserved (the studied populations of TEs). The proposed model is composed, in both of its variations (the one for conserved and the other for non-conserved elements), of biologically plausible molecular events and is based on a previous model formulated in the framework of aggregative dynamics^18^.

In Athanasopoulou *et al.* the entropic scaling analysis of the considered TE chromosomal distributions is accompanied by a box-counting study throughout^17^. Box-counting verified the appearance of fractality and self-similarity extending to several orders of magnitude in most cases where the aforementioned linear entropic scaling in semi-logarithmic scale was observed. In an older work of our group studying only chromosomal region (parts of chromosomes) where annotation about protein coding was available at the time, box-counting revealed indications of fractality in the distributions of genes^19^.

In references 1-3 and in numerous other later works, several research groups investigated aspects of genomic / nuclear structure at several length scales and levels of organisation. Their converging results indicate that long-range order, correlations extending at several length scales and fractality are ubiquitous in the nucleotide juxtaposition and in the distribution of functional elements or compositional inhomogeneities in the genome. In a study of Lieberman-Aiden et al.^20^ the organization of the eukaryotic nucleus according to the ‘fractal globule’ model has been proposed, through the combination of novel experimental and computational techniques. In this case, the fractal pattern has been revealed because the contact probability as a function of genomic distance across the genome shows a power law scaling at an important range of lengths. It is beyond the scope of the present article to exhaustively review this rapidly growing domain of research. In numerous works, Shannon entropy, fractality and related concepts (lacunarity, succolarity) are employed in order to describe genomic structure and to derive information which can be used from understanding functional and evolutionary aspects of genomic organization up to medical and diagnostic purposes. To mention a few, Cattani and Pierro (see ^21^ and references given therein) present a fractality and Shannon entropy analysis of whole chromosomes at the level of nucleotide distributions, deriving results which demonstrate the complementarity of the used methodologies. This study also suggests a framework applicable to the classification of DNA sequences. At a different level, evidence of fractality could be searched and quantified by means of an analysis of digitalized microscopic images of chromatin. Metze, in a comprehensive review^22^, describes how changes in the fractal dimension derived from image analysis can be applied in cancer prognosis. It has to be noted that such techniques are also applicable outside the chromatin research, as fractality appears in a multiplicity of microscopic and middle scale biological patterns with crucial roles in the understanding of the micro-anatomy of relevant tissues; see e.g. the work of Pantic and co-workers^23^ where fractality analysis of digital micrographs succeeds to systematically distinguish between histologically similar brain regions.

In the present work we focus on the study of several collections of Conserved Noncoding Elements (CNEs) using entropic scaling analysis and box-counting. The genesis and evolutionary dynamics of conserved noncoding elements remain an enigma. It has been calculated that approximately 5.5% of the human genome is under selective constraint; of that, 1.5% is believed to encode for polypeptide chains, 3.5% is assigned known regulatory functions, while there is little evidence suggesting possible roles for the remaining part^24^. The discovery of 481 ultraconserved elements (UCEs) of more than 200bp in length that are identical among human, mouse and rat genomes paved the way for a series of efforts with the aim to identify long sequences showing extreme levels of conservation^25^. Roughly 25% of those UCEs fall within known protein-coding sequences. Since the identification of UCEs, researchers have focused on identifying conserved elements based on (i) lower thresholds of sequence similarity over whole genome alignments of two or more organisms, (ii) several thresholds of minimal length of conserved sequence, and (iii) the filtering of elements located inside exonic sequences^26,27^. Throughout this article, we use the term CNE(s) for Conserved Noncoding Elements to refer to all such elements despite their specific characterization as UCNEs, CNEs, etc in the related literature. We use a particular name only whenever we want to refer to a specific class of elements.

It is believed that gene deserts are enriched in CNEs^28,29^ while, in mammalian genomes, a large number of those elements is often located at such distances from the closest genes that exceed in some cases 2 Mb, which is the limit for any known cis regulatory element^30,31^. Little could be conjectured concerning what those distant CNEs actually perform in the cell; there is evidence, however, showing that they form an essential part of Gene Regulatory Blocks (GRBs) and that they could synergistically function alongside with their target genes^32,33^. Furthermore, the literature suggests that CNEs are selectively constrained and not mutational cold spots^34^.

Since it is generally believed that sequence conservation across genomes is a key indication of functional relevance, the study of the sequence – specific characteristics of different classes of CNEs, as well as the mechanisms that might have led to their genesis, would be of paramount importance in an effort to crack the so far enigmatic regulatory code of our genome.

In a recent study of our group^7^, the size distribution of the inter-CNE distances of a variety of CNE collections has been investigated and power-law-like distributions have been found in most cases. This means that the abundance of inter-CNE spacers depends linearly on the spacers’ length in double-log scale, and this is found to occur for a range of spacers values often higher that two decimal orders of magnitude and in some cases exceeding the three orders. In the study presented herein we include entropic scaling and box-counting of four additional data sets not present in Polychronopoulos *et al*.^7^. For completeness, we include plots for the complementary cumulative size distribution of the inter-CNE distances for these CNE collections not studied earlier (see Supplementary Data File). We also refer the interested reader to this recent work for further details about aspects of CNE biology and several conjectures about their role and organization in the vertebrate genome.

## METHODS

### Box-counting method for the determination of fractality

Box-counting is widely used for assessing the fractality of symbol sequences and of other types of discrete datasets^35,36^. Here we describe an one-dimension implementation of this method. We cover the chromosome with one dimensional “boxes” of length *δ*. The number of boxes overlapping CNEs is assumed to be the chromosomal length L(*δ*) occupied by CNEs. In a fractal structure the length measured in this way does not reach a fixed value as *δ* decreases^36^. This length scales as: L(*δ*) ~ *δ*^D^. The exponent D is the negative fractal dimension D_f_ of the fractal pattern we consider. The plots depicting how L(*δ*) scales as a function of *δ* are shown in log-log scale. The slope of the linear part of the curve and the extent of the linearity are both informative for the characterization of the fractal pattern. Condition of existence of fractality is to obtain a value of D_f_ below 0.9 and a linear extent (*F*) exceeding, let us say, one order of magnitude. The limits of the linear region may be seen as the lower and upper cutoffs of fractality, and they determine the range of (statistical) self-similarity for the considered spatial structure. In order to obtain results independent of the choice of the starting point for the box-counting, for each value of *δ*, we compute L(*δ*) ten times, using a frame shift equal to 1/40 of the length of the sequence, and then we average.

In all chromosomal distributions of CNEs studied herein, two or three linearity regions are encountered. One is always found in the low-length region and is due to the (short relatively to inter-CNE distances) lengths of individual CNEs. The other one or two linear segments are located in the region of high length values and are due to the existence of lengthy spacers. The slope of these segments may deviate from -1 significantly, if fractality is present.

### Block entropy scaling

Let us consider a symbol sequence whose length is N, with symbols belonging to the binary alphabet {0, 1}. Let *p_n_(A_1_,…, A_n_)* be the probability for a block or *n*-word *(A_1_,…, A_n_)* to be present in the sequence. The definition of ‘block entropy’ or ‘Shannon-like entropy’ for *n*-words is (see reference 13):

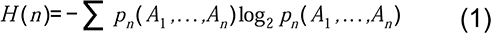

The most common interpretation for the entropy *H(n)* is that it represents a measure of the mean uncertainty in the prediction of an *n*-word. References 13-15 provide a survey of the main properties of block entropy and of other similar quantities. Only a summary of principal results of direct relevance to our study will be given here, while a more detailed analysis is included in reference 12.

Two standard ways exist for the reading of symbol sequence and for the extraction of the probability distribution of *n*-words: ‘gliding’ and ‘lumping’. In the present work, ‘lumping’ has been used throughout. During this mode of reading, we do not proceed to the exhaustive reading of all possible words of length *n* (this is the method of gliding). Only *n*-words sampled with a constant step equal to *n* are considered. This is equivalent to say that after reading the initial *n*-word of the sequence, the next counted *n*-word is the one starting at *n*+1 and so on up to the end of the sequence.

Block entropy presents scaling properties which may serve the purpose of classification of the symbol sequences according to their long-range properties. Ebeling and Nicolis^14^ have proposed the form of equation (2) for the scaling of *H(n)*:

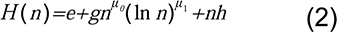

in the case of symbolic sequences which are the outcome of a non-linear dynamics, especially in cases of language-like processes^14–16^. In the very important case of the Feigenbaum attractor of the logistic map and for *n=2^k^* (k=2, 3, 4 …), Grassberger^13^, see also^37,38^, has shown that, the scaling is of the form:

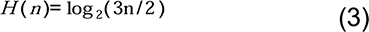

if the sequence is read by gliding. In this system, linearity in semi-logarithmic plot holds, which in terms of equation (2) corresponds to: *^g^* ^≠ 0^, *^h^* ^= 0^, *^μ^*^0= 0^, *^μ^*^1> 0^, ^39^. This form of entropic scaling is assumed to be valid for a wide range of fractal symbol-sequences. As a consequence, the linearity between *H(n)* and log*n* is attributed to the scale-free structure of these sequences which gives rise to long-range correlations. In a previous work^12^ we have verified this hypothesis for Cantor-like symbol-sequences (of both, deterministic and probabilistic type).

For every chromosomal data set we have produced a surrogate random sequence retaining its ‘0’/’1’ composition but without any trace of the initial internal structure left. The entropic scaling (*H(n) vs. n*) curve of the initial chromosome and that of the surrogate one are presented within the same plot. Random surrogates are also used along with model generated sequences.

The extent (E) of linearity in semi-logarithmic scale is used as a quantifier of fractality of a chromosomal distribution of CNEs. When two linear segment exist, we symbolize with E^*^ the sum of their lengths. We introduce and use heuristically as an estimator of the degree of internal structure of a sequence the ratio R of the entropy value of the surrogate sequence over the entropy value of the studied (genomic or simulated) sequence. R is computed for the value of *n* at which the surrogate sequence reaches its maximum entropy. Beyond this value its entropy starts decreasing due to finite size effects. High values of R denote a high degree of order of the studied sequence.

### Datasets

We consider published CNE datasets, derived from whole genome alignments of the human with other genomes, ranging from elephant shark to mammals:

i. 82,335 ‘Mammalian’ CNEs, mapped on the human genome (hg17)^28^.
ii. 16,575 ‘Amniotic’ CNEs, mapped on the human genome (hg17)^28^.
iii. 3,124 ‘Human – Fugu’ CNEs mapped on the human genome (hg17)^40^.
iv. 4,782 ‘Human – Elephant Shark’ CNEs mapped on the human genome (hg17)^41^.
v. 2,614 ‘Rodent’ extremely constrained sequences mapped on hg17, some of which act as developmental enhancers^42^.
vi. 4,386 ‘UCNEs’ or ‘CNEs 95-100’ (Ultraconserved Noncoding Elements, longer than 200bp) mapped on the human genome (hg19) that display sequence similarity in the range of 95% - 100% between human and chicken whole genome alignments^33^. These elements represent the most constrained subset of sequences, while the next four sets have been described in a recent study using less stringent thresholds of sequence similarity^43^.
vii. 4,635 ‘CNEs 90-95’ (longer than 200bp) mapped on the human genome (hg19) that display sequence similarity in the range of 90% - 95% between human and chicken whole genome alignments^43^.
viii. 5,860 ‘CNEs 85-90’ (longer than 200bp) mapped on the human genome (hg19) that display sequence similarity in the range of 85% - 90% between human and chicken whole genome alignments^43^.
ix. 7,615 ‘CNEs 80-85’ (longer than 200bp) mapped on the human genome (hg19) that display sequence similarity in the range of 80% - 85% between human and chicken whole genome alignments^43^.
x. 10,010 ‘CNEs 75-80’ (longer than 200bp) mapped on the human genome (hg19) that display sequence similarity in the range of 75% - 80% between human and chicken whole genome alignments^43^.

For additional details about the used data sets and the subsequent treatment see Supplementary Data File. The suite of utilities BEDTools has been used for the genomic analyses^44^.

### Masking of genic regions

We have masked the entire sequence stretches annotated as genic in the human genome. Furthermore, we have masked flanks surrounding every gene: 5 kb at both 5’ and 3’ ends, so that we cover regulatory elements the position of which might be principally determined by the spatial organization of the gene under regulation. The region located upstream of transcription start sites is usually particularly enriched in such regulatory sequences. Extended symmetric flanks of 50 kb or 100 kb have also been masked, where possible, in a similar way (see “Results” section). We use custom scripts and BEDTools in order to perform the masking. Note that by masking, we mean shadowing all those elements that overlap genes and flanking regions and not removing the genes themselves, as the latter would affect the studied genomic landscape. For genomic coordinate data of masked genes, see Supplementary Data File.

We do not proceed to the masking of other specific classes of sequences, e.g. transposable elements, that have been shown to form fractal-like patterns within the genome, because there is no evidence suggesting CNE – TE functional inter-play or systematic co-localization. Only a small proportion of TEs is reported to have been exapted to the role of a CNE, but those are too few to influence and alter the entire CNE distribution^45^.

### The “Segmental-duplication / CNE elimination” model

In previous works we formulated a version of the aforementioned evolutionary model describing genomic dynamics relevant to the evolution of Protein Coding Segments (PCSs) or of CNEs, based on quite general prerequisites, mainly related to the conserved character of these genomic elements and the underlying genomic dynamics^4,7^. This model, through computer simulations, generates fractal-like patterns, like the ones observed in real chromosomes. Ηere we only briefly describe the structure of the model while we refer the reader to these previous works for a detailed description of its biological background. The “Segmental-duplication / CNE elimination” model dynamics builds upon models for the explanation of fractality in aggregation patterns in physicochemical systems^18^. Our model takes into account the one-dimensional topology of DNA and includes molecular events known to have occurred in genome dynamics over the course of evolutionary time. Note that we have also introduced elsewhere a different model which focuses on the genomic dynamics of mostly non conserved elements, like TEs, and accounts for the particular distributional characteristics of these elements in genomes.

Occasional loss of function of some CNEs and their subsequent degradation^46^, whole genome duplications and regularly occurring segmental duplications^47–51^ together with other types of genome dynamics (e.g. insertions of transposable elements and other parasite sequences) can be integrated in the “Segmental-duplication / CNE elimination” model. Its propensity to generate fractality, as evaluated through box-counting, and the aforementioned entropic scaling, both observed in chromosomal distributions, is testable by means of computer simulations. The genomic events included are:

i. Segmental duplications of chromosomal regions.
ii. Eliminations at random of a number of CNEs which is lower or equal to the number of those ones that get duplicated.
iii. Additional eliminations of CNEs that have not undergone duplication.
iv. Insertions of sequences that expand the chromosomal length (for example insertion of transposable elements, microsatellite expansions or intruding retroviruses etc).
v. Deletions of sequence regions (assuming that they are under weak or no purifying selection).

Characteristic cases of simulations are shown in the last two figures. Note that we have chosen to include here the same examples of simulations which are included in the aforementioned work where the chromosomal distributions of CNEs had been studied examining the form of the cumulative distribution of inter-CNE distances^7^. Initially, 1000 markers (representing CNEs) are randomly dispersed in a sequence 2 Mbp long. The box-counting and entropic scaling plots corresponding to the initial random CNE distribution may be found in the Supplementary Data File. The penultimate figure depicts snapshots of the emerging distributional pattern through time. Plots are generated every 50 segmental duplication events. The length of the region involved in each segmental duplication (SD) event is obtained from a uniform distribution with maximum the 5% of the actual length of the simulated sequence. A number of CNEs equal to 90% of the number of the duplicated CNEs (denoted as: fr = 90%) is eliminated after each SD event in these numerical simulations. No further eliminations of CNEs that have not undergone duplication are taken into consideration here. In the upper half of the last figure, plots for numerical experiments, where fr takes the values 80% and 100%, are shown. These plots must be evaluated in comparison to parts e & f of the penultimate figure, as for all of them, 150 SD events have been simulated. Next, in the lower half of the last figure, the plots for numerical experiments are presented, where the number of SD events is the same, but additionally, we consider eliminations of non-duplicated CNEs (one or two after each SD event). These plots have to be evaluated in comparison to parts e & f of the penultimate figure again. In the last table, the quantitative aspects of these numerical experiments are summarized, while in the “Discussion” section, conclusions about the convergence between genomic and simulated chromosomal distributions of CNE are drawn.

## RESULTS

Entropic scaling and box-counting are performed for the CNE sets which are described in the “Methods” section for all chromosomes of the human genome. In each case, chromosomes are treated as symbol-sequences by replacing nucleotides belonging to the considered CNE set with ‘1’s while replacing with ‘0’s the rest of the nucleotides. The overall result of the present study is that almost all CNE collections we have examined herein present fractality, often highly developed, in some cases above 3 orders of magnitude, as measured through box-counting. Their entropic scaling reveals that most of them exhibit considerable linearity in semi-logarithmic scale and a ratio R well above unity. Although R does not represent a measure of fractality, it indicates the extent to which a given genomic sequence has reduced its block entropy (thus, it might be seen as more organized) with respect to its random surrogate. Block entropy is expected to be strongly reduced when fractality is developed, and inspection of the last table, where metrics of model-generated sequences are presented, corroborates this view.

### Fractality measured by box-counting and entropic scaling in the distribution of various CNE collections in entire chromosomes

Figure 1 shows eight examples of box-counting plots for different CNE datasets in entire human chromosomes. Quantification of fractality is effected by means of the extent F of linear segments in log-log scale and the associated slope which equals the negative fractal dimension D. The convention we follow here is that D values around 1 (+/− 0.1) denote lack of fractality, while fractal-like geometry is present when D falls lower than 0.9. We always observe lack of fractality in the linear segment for the low length regions (at the left side of the plots). When we replace the CNE in each chromosome with a single ‘1’ symbol, the low length linear segment disappears while the rest of the plot remains unaffected. This indicates that this linearity relates to the length variation of the CNEs which as expected is short ranged, and thus, shows no trace of fractality. Such plots are not presented here, as a similar analysis is presented in the case of our study for the chromosomal distribution of TEs. On the other hand, at the high length region, linearity with absolute slope clearly lower than unity is often observed. In several cases, this linearity is divided into two regions, which indicates that the scaling features of the CNE chromosomal distribution are different at different length scales. This displays an analogy with the similar observation (see Figure 2 and Table 1) that segmented linearity is also exhibited in the entropic scaling plot. This feature is compatible with the non-universality of the model we propose (see in the “Methods” and the “Discussion”) where also a unique slope is not generated.

**Figure 1:**
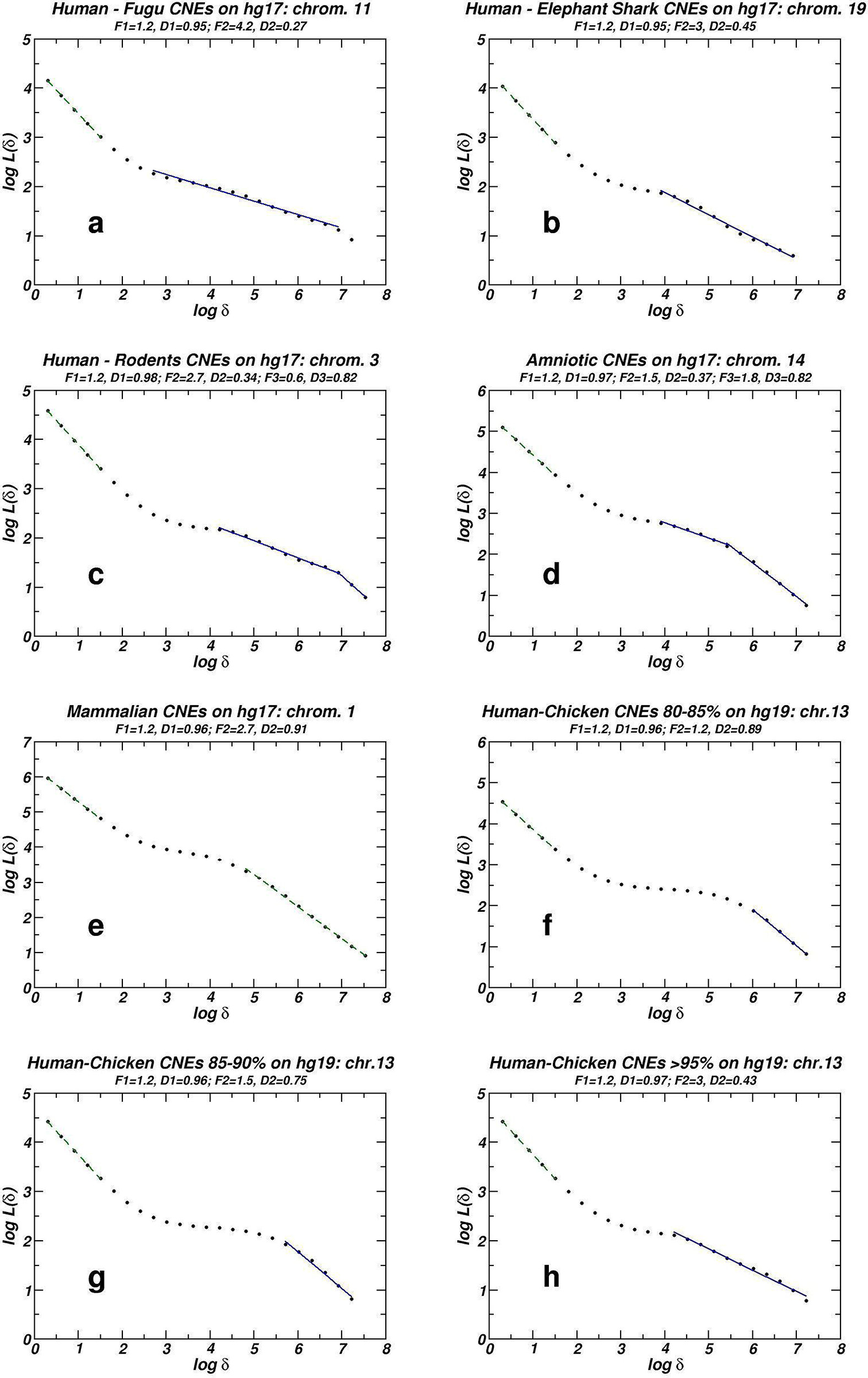
Examples of whole chromosome box-counting plots for eight different CNE datasets. Linear segments are generated by linear regression. Solid and dashed lines stand for presence and absence of fractality respectively.

**TABLE 1^*^.**

| Conserved Noncoding Elements datasets. | *Box-counting plots* |  | *Entropic scaling plots* |  |  |
| --- | --- | --- | --- | --- | --- |
| | Average<sup>#</sup> Fractality-related Linearity (FrL)<sup>&</sup>: $F_2 + F_3$ | Average of the 5 highest FrL values | Average<sup>#</sup> Linearity $E$ or $E^*$ (1 or 2 linear<sup>§</sup> segments) | Average of the 5 highest linearity values | Average<sup>#</sup> of $R$ values<sup>§</sup> |
| 'Mammalian' CNEs | 2.05 | 2.10 | 0.78 | 1.10 | 1.48 |
| 'Amniotic' CNEs | 2.39 | 3.06 | 1.45 | 1.93 | 1.88 |
| 'Rodent' (Homo – Mouse – Rat) CNEs | 2.57 | 2.95 | 1.60 | 1.92 | 1.99 |
| 'Homo – Elephant Shark' CNEs | 2.76 | 3.07 | 1.68 | 2.35 | 2.50 |
| 'Homo – Fugu' CNEs | 3.20 | 3.60 | 1.98 | 2.39 | 2.81 |
| 'Homo – Chicken' CNEs Seq. similarity 75 - 80% | *No fractality found* |  | 0.85 | 1.00 | 1.25 |
| 'Homo – Chicken' CNEs Seq. similarity 80 - 85% | 1.64 | 1.95 | 1.04 | 1.21 | 1.31 |
| 'Homo – Chicken' CNEs Seq. similarity 85 - 90% | 2.45 | 2.94 | 1.11 | 1.36 | 1.44 |
| 'Homo-Chicken' CNEs Seq. similarity 90 - 95% | 2.81 | 3.18 | 1.12 | 1.64 | 1.63 |
| 'Homo-Chicken' CNEs Seq. similarity > 95%, also called 'UCNEs' | 2.81 | 3.12 | 1.49 | 2.04 | 2.47 |

* Detailed tables of all F1, F2, F3 for box-counting plots and E or E* along with R for entropic scaling plots for all studied CNE sets are given in Supplementary Spreadsheet and the corresponding plots in Supplementary Data File.
\# Averages are calculated for all human chromosomes where the CNE instances of a given CNE collection are more than 100. & Using the box-counting method linearity in double-log scale is considered to express fractality if its absolute slope is less than 0.9. When two such linear segments are found, the overall linearity is denoted as Fractality-related Linearity (FrL).
§ In entropic scaling plots linearity is studied in semi-log scale and is denoted by E. When two linear segments are found, the overall linearity is denoted by E*.
$ In entropic scaling plots deviation from randomness is also measured by means of the ratio R of the entropies of surrogate over genomic sets measured for the word lengh where the maximum in the surrogate sequence curve is observed (for details see in the “Methods”).

In Figure 2 we present eight examples of *H(n)* plotted versus *n* (the block- or word-length) for several CNE collections in different human chromosomes. In each plot, the curve for the genomic curve is accompanied by the curve of a surrogate sequence of length equal to the length of the studied chromosome and the same number of CNEs positioned at random. The degree of entropic scaling is measured by the extent E of the linearity in semi-logarithmic scale as well as the ratio R of the entropy values of the surrogate and the genomic curve computed at the maximum of the surrogate curve (see in the “Methods” section for details).

**Figure 2:**
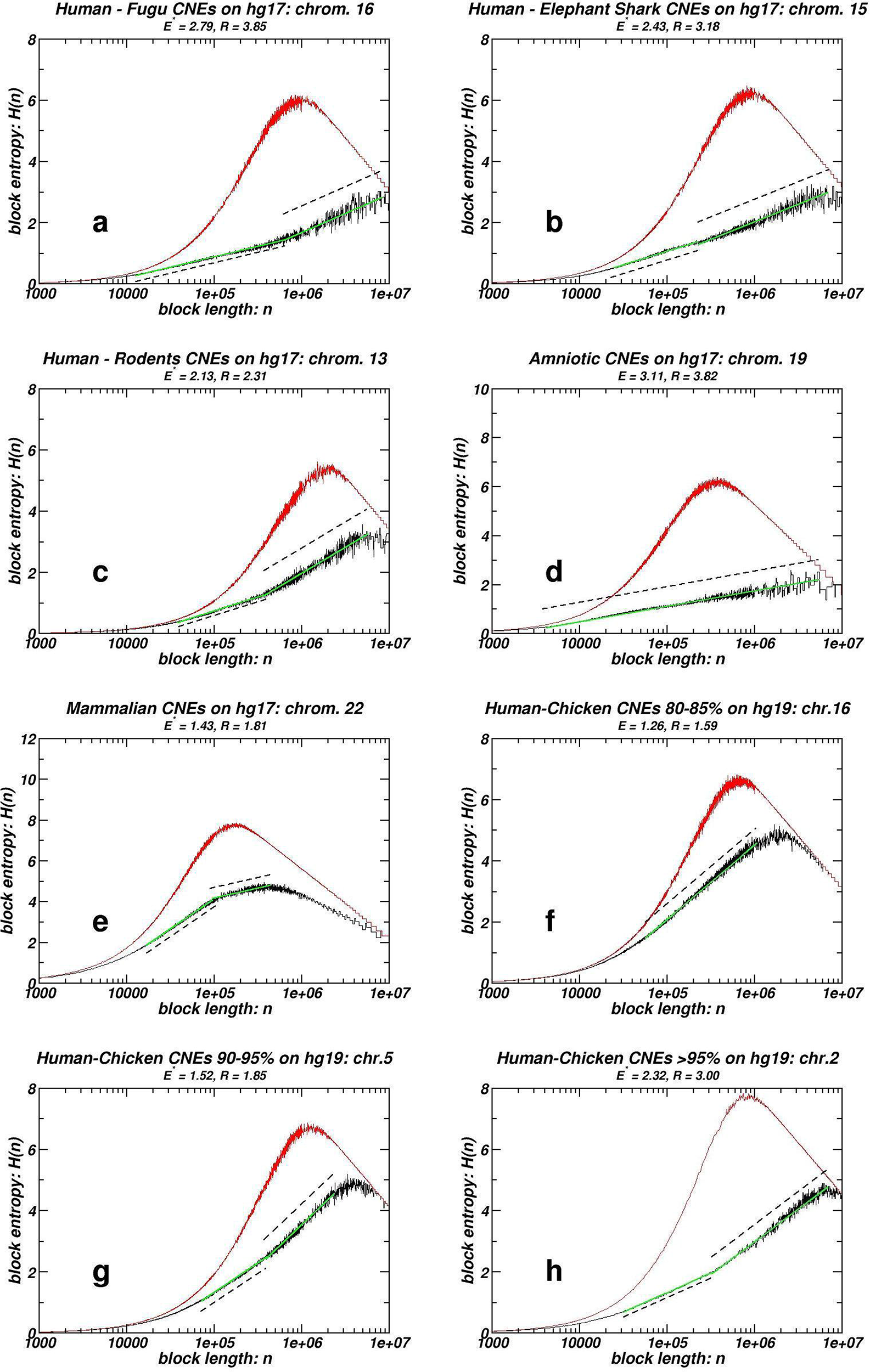
Examples of whole chromosome block entropy H(n) plots for eight different CNE datasets. Random surrogates are also included. Genomic sequence is shown in black and the random surrogate is shown in red. Dashed linear segments are parallel to the linear regression green line.

In Table 1, average values of all human chromosomes for each CNE dataset of: **(i)** The Fractality-related Linearity (FrL) which is the sum of the box-counting linear segments corresponding to fractal dimension lower than 0.9; **(ii)** Linearity E (or E^*^ if more than one linear segment is found); and **(iii)** The ratio R. Additionally, averages of the five highest values for FrL and E are also provided in Table 1 as additional quantifiers of the box-counting and entropic scaling fractality for each CNE collection.

In the Spreadsheet within the Supplementary Data File, metrics for all individual chromosomes for all CNE datasets are provided. Simple inspection reveals that these metrics differ for the several collections. This variation lies in accordance with the proposed model, as discussed below.

### Fractality measured by box-counting and entropic scaling in the distribution of CNE collections in gene-masked chromosomes

As briefly presented in the “Introduction”, fractality revealed through the box-counting and entropic scaling approaches is evidenced in the chromosomal distribution of protein-coding segments^12,19^. Published studies also suggest that some CNEs are spatially linked with genes that mostly encode for transcription factors and regulators of development (collectively referred as trans-dev genes)^30,52^. To exclude the possibility that the fractality patterns reported herein stem from the analogous patterns followed by genes^12^, we mask all protein coding sequences as well as extended proximal regulatory regions, in the three of the studied datasets exhibiting the most pronounced fractality. In Figures 3 and 4 we show plots analogous to the ones presented in Figures 1 and 2, while in Table 2 average quantities for masked chromosome sets are provided, following the conventions set in Table 1. Flanks of 5 kb are used upstream and downstream of each gene, while occasionally extended symmetric masking of 50 and 100 kb has been employed. The corresponding plots are given in the Supplementary Data File. All the quantities for individual chromosomes are given in the Spreadsheet within the Supplementary Data File. The chromosomes that contain less than 100 CNE instances, a case often encountered after masking, are excluded from our study due to poor statistics. The observed linearities in log-log and semi-log scale for the box-counting and entropic scaling plots respectively, as well as R values in entropic plots, are not only preserved but slightly increased in all studied cases. The comparisons are made with averages of the corresponding values of the same unmasked chromosomes (see numbers in parentheses in Table 2). This indicates that even if we exclude from our study those CNEs that might be associated somehow with genes due to their spatial co-occurence, the remaining CNEs still follow the chromosomal distribution that appears in the cases of the non-gene-masked genomes. Even if extended masking is applied, fractality is still evident.

**Figure 3:**
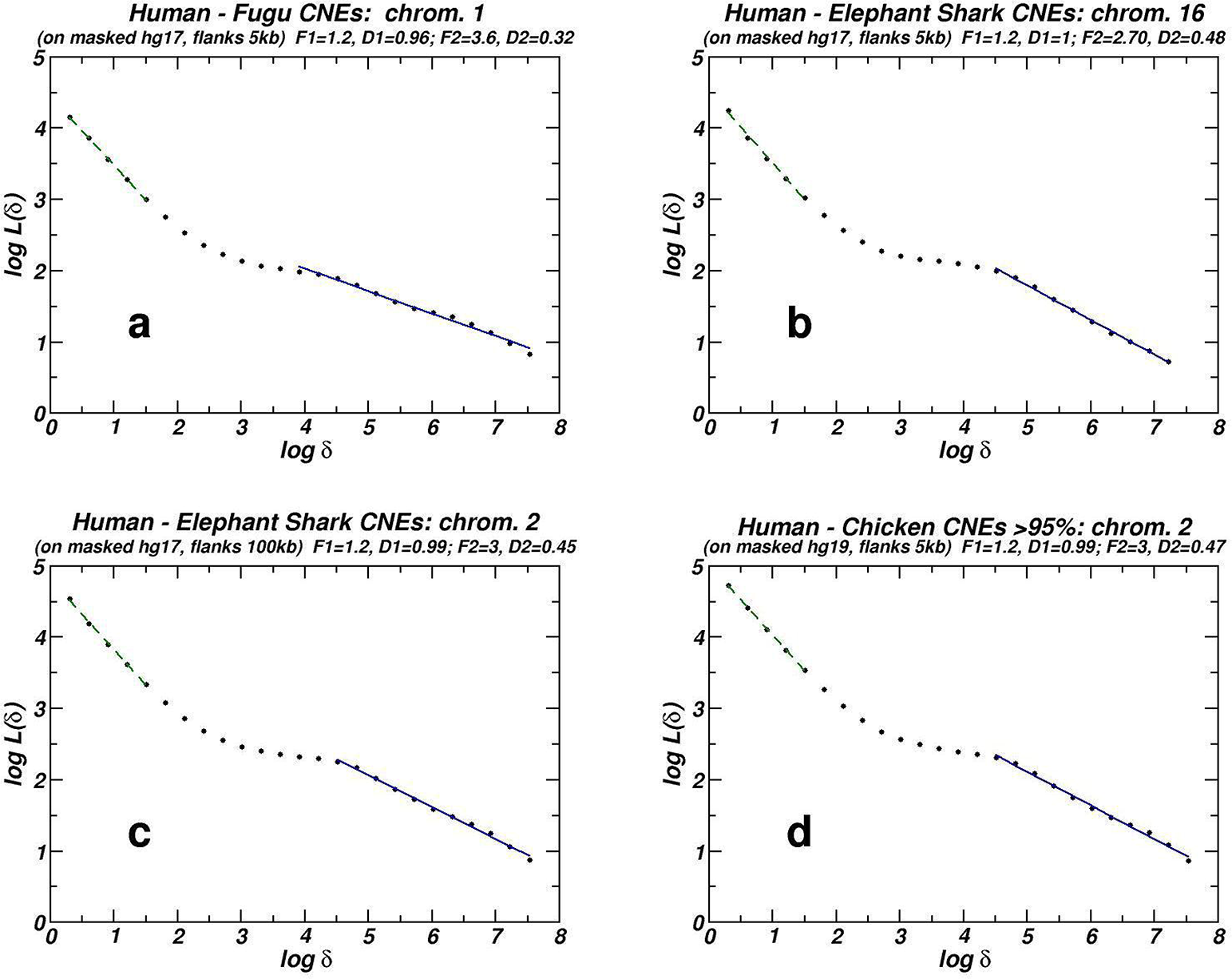
Examples of whole chromosome box-counting plots for CNE datasets after masking of genes and symmetric 5’ and 3’ flanks of 5 kb or 100 kb each. Plotting conventions are as in Figure 1.

**Figure 4:**
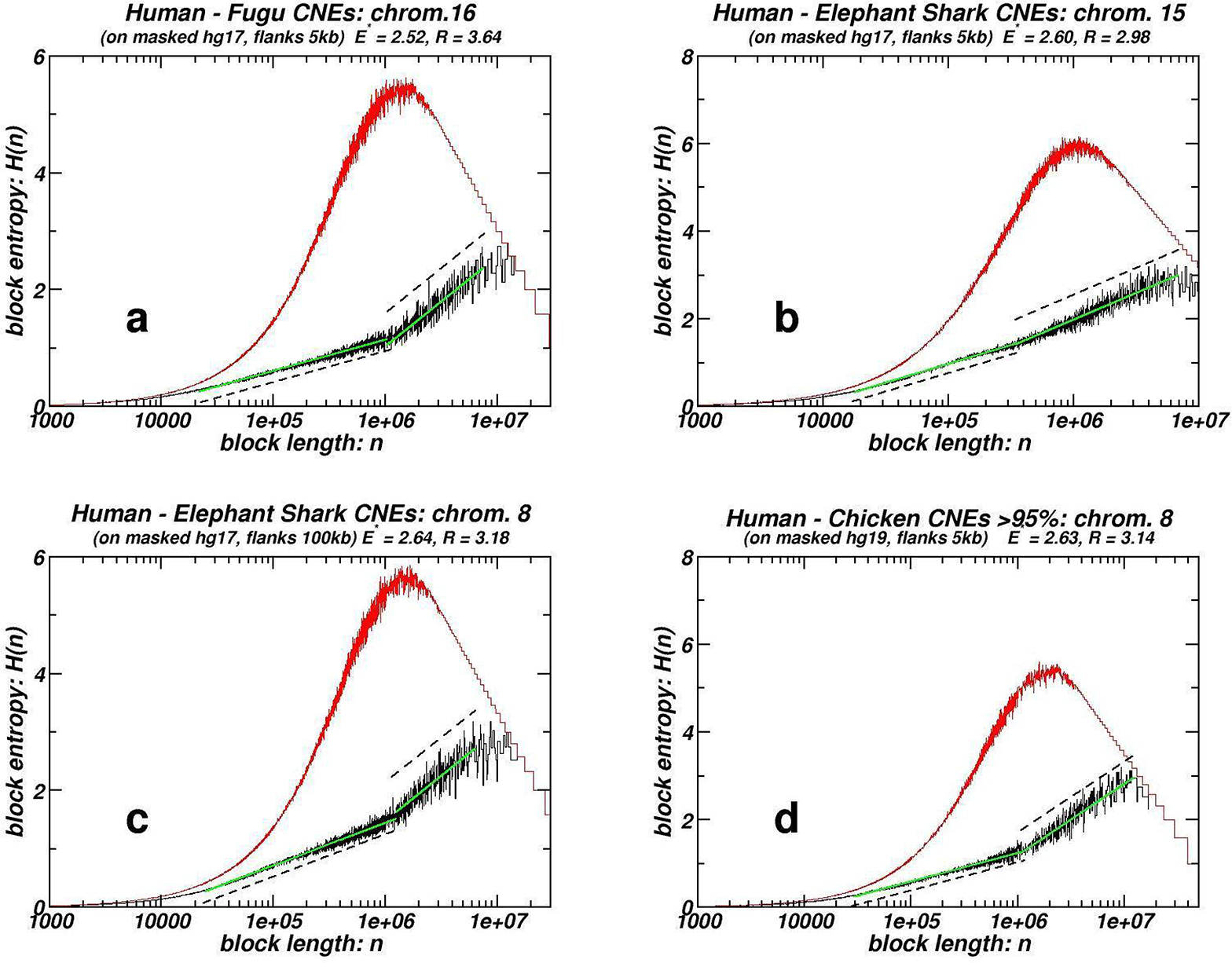
Examples of whole chromosome block entropy H(n) plots for CNE datasets after masking of genes and symmetric 5’ and 3’ flanks of 5 kb or 100 kb each. Plotting conventions are as in Figure 2.

**TABLE 2^*^.**
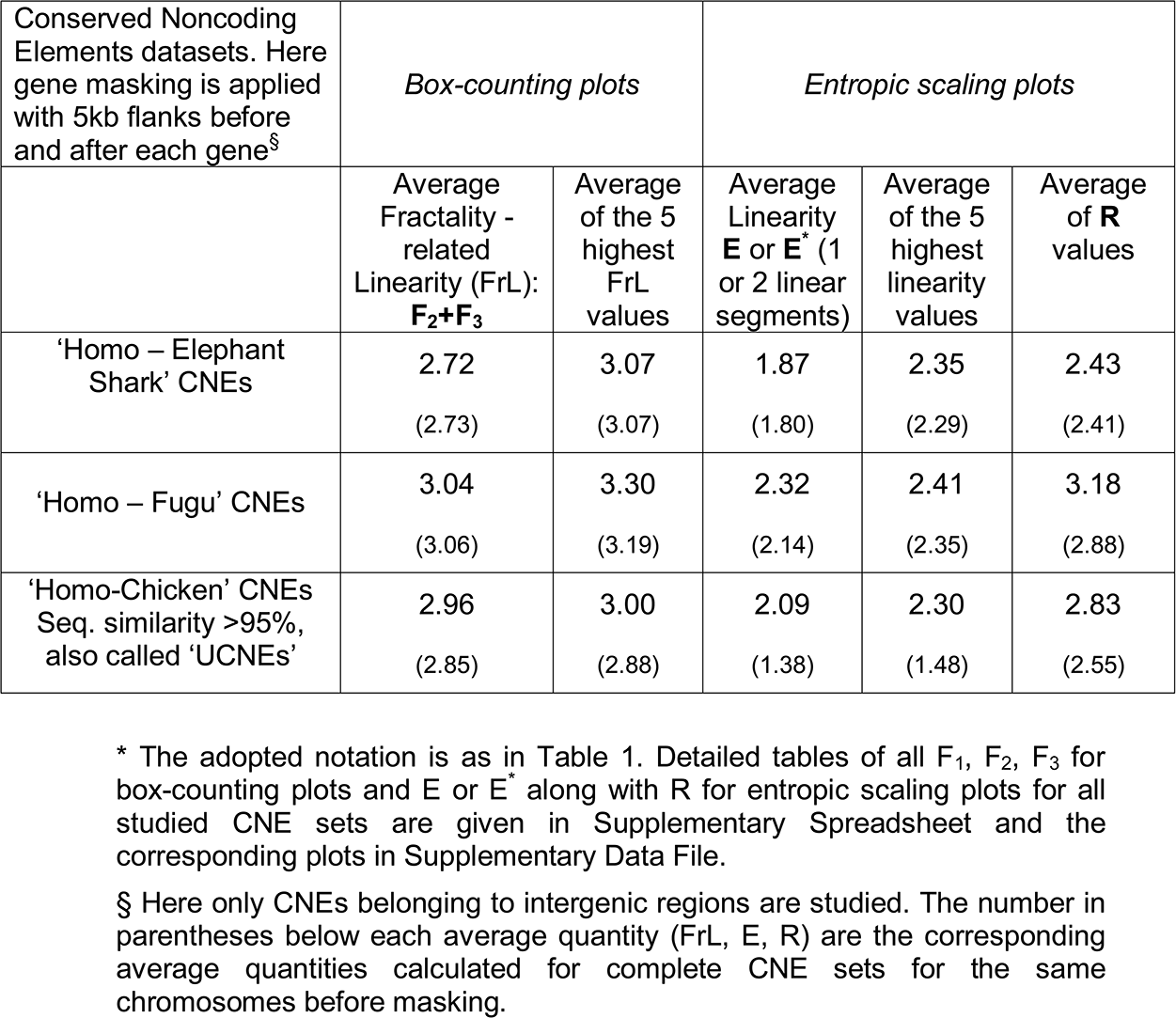

### Box-counting and entropic scaling of CNE distributions derived by the “Segmental-duplication / CNE elimination” model

In Figures 5 & 6, plots for numerical experiments of the proposed model are shown. The corresponding metrics are presented in Table 3. Three different aspects of the model are studied that correspond to key variables of the underlying molecular dynamics. In Figure 5 and in the first part of Table 3, the effect of allowing Segmental Duplications (SD), a common phenomenon that is known to have shaped the human genome^47–51^, is monitored as successive snapshots are taken, each after 50 additional SDs. As described in the “Methods”, elimination of most of the duplicated CNEs and, occasionally, loss of non-duplicated ones are incorporated in the model. These molecular events are introduced into the model through the percentage of duplicated CNE which are lost (denoted by fr) and a number of lost non-duplicated CNEs between consecutive Segmental Duplication events. In the numerical experiment depicted in Figure 5, we set fr equal to 90% and no loss of non-duplicated CNEs is allowed. The plots corresponding to the random initial condition of the numerical experiment are shown in Supplementary Data File. As more and more events of SDs occur, we observe a progressive increase of all metrics reflecting fractality of the final CNE distribution in the artificial simulated chromosome. If they occur through time in a constant or variable rate, the proposed model predicts that the more ancient a dataset of studied CNEs is, the more pronounced the fractal properties of its chromosomal distribution will be. Bursts of duplication activity or whole genome duplications events^51,53^ are expected to contribute to the emergence of fractality.

**TABLE 3^*^.**
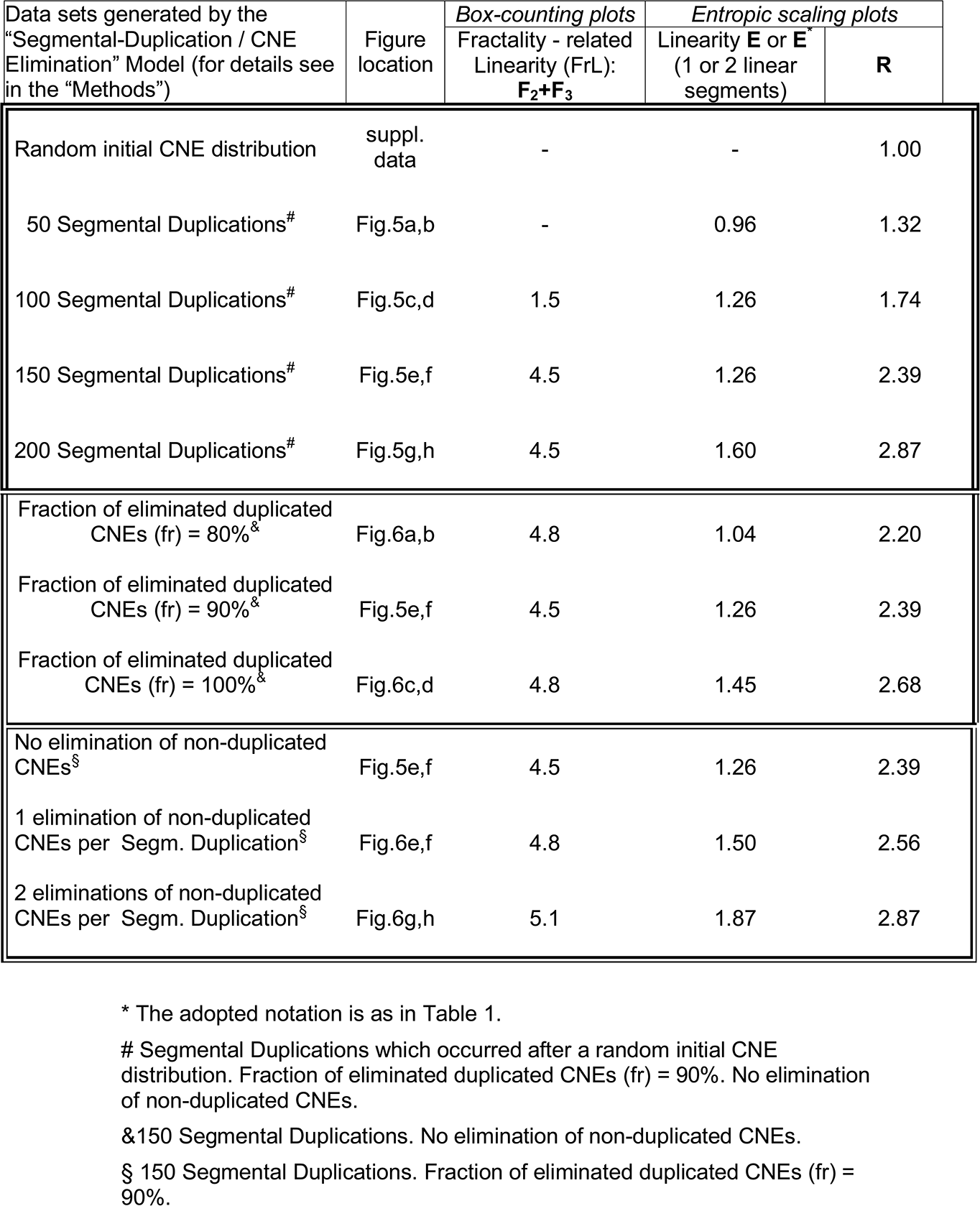

| Data sets generated by the<br>“Segmental-Duplication / CNE<br>Elimination” Model (for details see<br>in the “Methods”) | Figure<br>location | Box-counting plots | Entropic scaling plots |  |
| --- | --- | --- | --- | --- |
|  |  | Fractality - related<br>Linearity (FrL):<br>**F<sub>2</sub>+F<sub>3</sub>** | Linearity **E** or **E\***<br>(1 or 2 linear<br>segments) | **R** |
| Random initial CNE distribution | suppl.<br>data | - | - | 1.00 |
| 50 Segmental Duplications<sup>#</sup> | Fig.5a,b | - | 0.96 | 1.32 |
| 100 Segmental Duplications<sup>#</sup> | Fig.5c,d | 1.5 | 1.26 | 1.74 |
| 150 Segmental Duplications<sup>#</sup> | Fig.5e,f | 4.5 | 1.26 | 2.39 |
| 200 Segmental Duplications<sup>#</sup> | Fig.5g,h | 4.5 | 1.60 | 2.87 |
| Fraction of eliminated duplicated<br>CNEs (fr) = 80%<sup>&</sup> | Fig.6a,b | 4.8 | 1.04 | 2.20 |
| Fraction of eliminated duplicated<br>CNEs (fr) = 90%<sup>&</sup> | Fig.5e,f | 4.5 | 1.26 | 2.39 |
| Fraction of eliminated duplicated<br>CNEs (fr) = 100%<sup>&</sup> | Fig.6c,d | 4.8 | 1.45 | 2.68 |
| No elimination of non-duplicated<br>CNEs<sup>§</sup> | Fig.5e,f | 4.5 | 1.26 | 2.39 |
| 1 elimination of non-duplicated<br>CNEs per Segm. Duplication<sup>§</sup> | Fig.6e,f | 4.8 | 1.50 | 2.56 |
| 2 eliminations of non-duplicated<br>CNEs per Segm. Duplication<sup>§</sup> | Fig.6g,h | 5.1 | 1.87 | 2.87 |

\* The adopted notation is as in Table 1.

**Figure 5:**
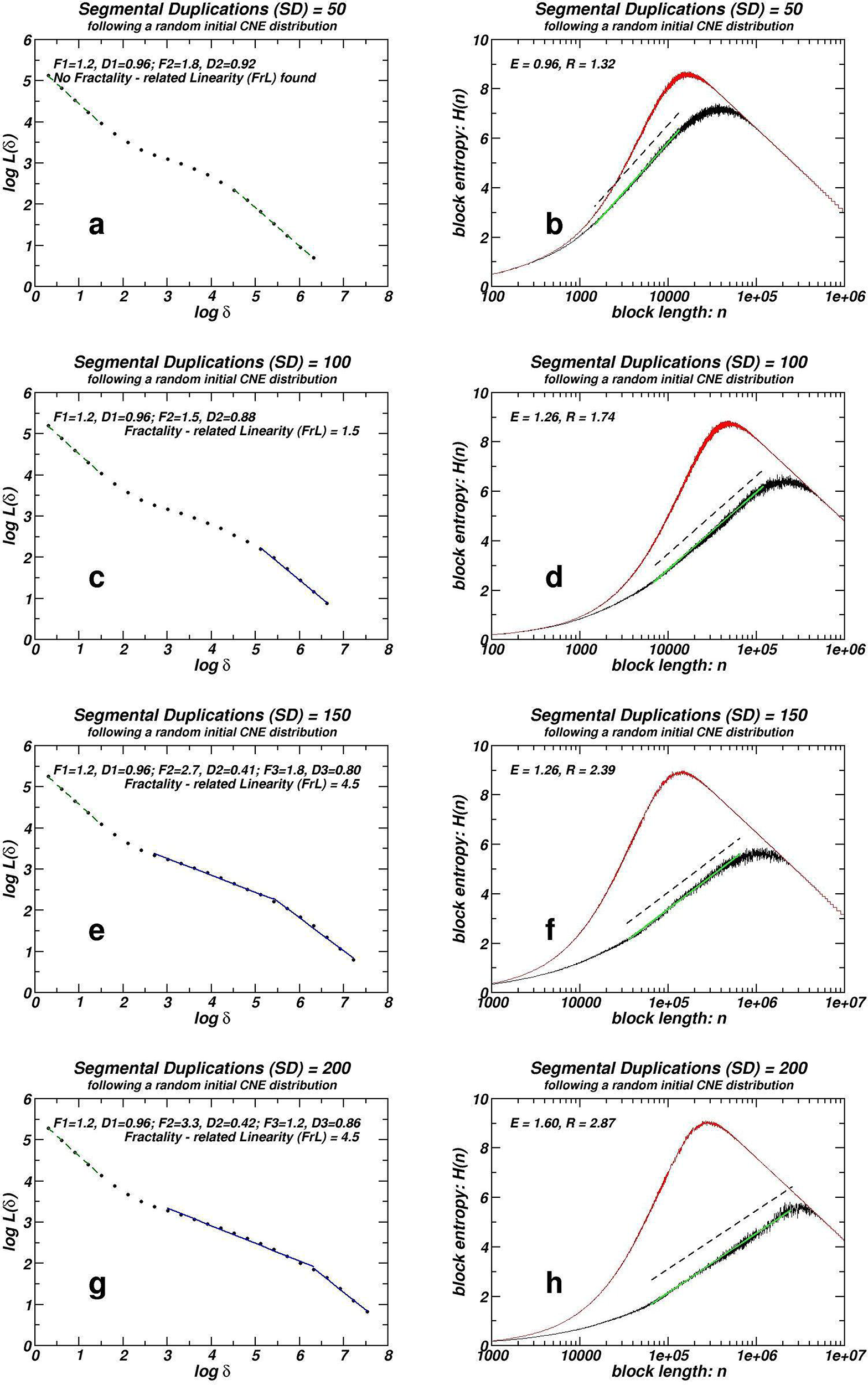
Box-counting and block entropy H(n) plots for numerical simulations according to The “Segmental-duplication / CNE elimination” model. The number of segmental duplications increases with a step of 50 SDs. In all cases, a number of CNEs equal to the 90% of the duplicated ones (denoted as fr = 90%) are eliminated after each segmental duplication. No eliminations of non-duplicated CNEs occur. Box-counting and entropic scaling plots corresponding to the initial random CNE distribution may be found in the Supplementary Data File.

**Figure 6:**
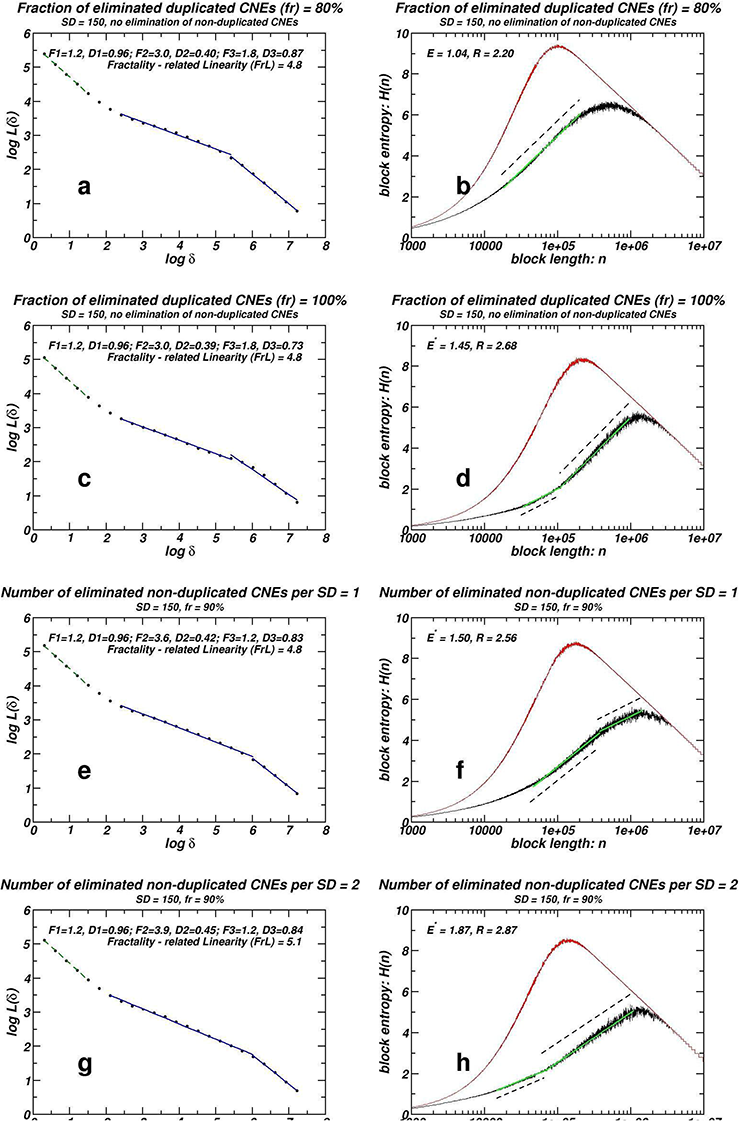
Box-counting and block entropy H(n) plots for numerical simulations according to The “Segmental-duplication / CNE elimination” model. The number of segmental duplications remains constant (SD = 150, cf. Figure 5e&f). In the upper half of the Figure the percentage of duplicated CNEs (fr) and in the lower half the number of non-duplicated CNEs which have been lost after each SD vary.

In Figure 6, the impact of the loss rate of duplicated (upper half) and non-duplicated CNEs (lower half) is monitored. In Table 3 (second and third part respectively) the corresponding quantities are given. In most cases, we observe increase of metrics indicating fractality, when loss of either duplicated or non-duplicated CNEs increases. Assuming that the proposed model describes the essential dynamics responsible for the emergence of fractality, the CNE datasets identified in taxa which have incurred the most CNE losses would exhibit the most well developed fractality.

As we will see in the following section, the predictions based on the simulations presented herein (Figures 5, 6, and Table 3) fit well with what is observed in genomic CNE datasets (Figures 1, 2, and Table 1).

## DISCUSSION

Fractality is widespread in the chromosomal distribution of CNEs identified using various threshold sets, between organisms of various evolutionary distances. We have to stress here that fractality as measured in different datasets does not indicate universality, as no unique fractal dimension is reached. This complies with the proposed model, where, also the obtained slopes vary, depending on the parameters, although in both genomic and model-derived sequences, extended linearity has been obtained in several cases. An analogous picture emerges in the study of power-law-like distributional patterns for many CNE datasets and for sequences generated by using the same model^7^.

In order to compare the degree of capturing self-similarity and fractality features in genomic sequences by means of inter-CNE distances statistics, entropic scaling and box-counting, we focus on the ‘mammalian’ dataset. CNEs belonging to this dataset have been exapted in a relatively recent time period, and thus they do not have adopted the self-similar pattern in all chromosomes. From reference 7 we deduce that, in 19 out of 22 human chromosomes (where such CNEs have been found), a power-law-like distribution pattern has been identified, although with only a moderate extent of the linear segment, if compared to all the other studied datasets. In the present work we find fractality by means of box-counting in only six chromosomes for the same dataset. The inclusion of entropic scaling in this comparison is not straightforward, as a moderate linear segment in semi-log scale and an R value just above unity are widespread and even intuitively they cannot be assigned to the occurrence of a clear fractal / self-similar pattern. If, however, we set a threshold of ¾ orders of magnitude extent of linearity (E > 0.75), we find only five chromosomes exceeding this value. If we set a threshold of 1.5 for the ratio R, we find again five chromosomes located above this threshold. Thus we might conclude that box-counting and entropic scaling constitute methodologies more stringent than the simple distance statistics (expressed in the form of power-law distributions between the distances of consecutive CNEs) for the evaluation of long-range order and emergence of fractality in symbol sequences.

The inspection of cases of gene-masked datasets (see Figures 3, 4 and Table 2) shows that the dynamics that shaped the fractal-like pattern is not a simple result of the coexisting distribution of protein-coding genes, notwithstanding that the two populations of genomic elements are expected to influence one another, although such interactions are not included in our simple model. The slight increase of fractality observed in the masked chromosomes is probably an indication that, when all the CNEs are examined together, the resulting distributions might reflect a superposition of two distinct dynamics. These different dynamics should express the differences in the evolution and thus in the genomic modalities of gene-uncorrelated CNEs and of genes (with which gene-proximal CNEs are spatially associated). This superposition of distributions with different spatial characteristics is expected to reduce the observed linearity extent in box-counting and entropic scaling plots in double logarithmic and semi-logarithmic scale respectively.

The proposed model predicts that more ancient CNE datasets are expected to exhibit more developed fractality compared to more recently exapted ones. Inspection of our findings as tabulated in Table 1 does support this prediction in all cases where a relation of age between several CNE datasets can be established. ‘Mammalian’ and ‘Amniotic’ CNE collections represent the most clear such case, as they were generated using the same parameters (minimal sequence length and similarity thresholds) by the same research team, while they were exapted in different evolutionary periods, with ‘Amniotic’ being older. Fractality as measured by box-counting and both linearity extent and the ratio R in the entropic scaling representation are higher in the ‘Amniotic’ than in the ‘Mammalian’ dataset. Comparison between ‘Rodent’ and ‘Homo – Elephant Shark’ datasets, which are also of clearly different evolutionary depth, again verifies this tendency.

In the list of the used datasets (see “Methods”), four recently identified CNE collections^43^ display sequence similarity between human and chicken whole genome alignments gradually increasing from 75% to 95%. They have been numbered datasets ***(vii) – (x)*** and their box-counting and entropic scaling properties are listed (see Table 1) along with dataset ***(vi)*** which consists of the most conserved ‘Homo – Chicken’ CNEs with similarity above 95% (and up to 100%) named UCNEs^33^. A gradual increase of all their fractality metrics is evidenced, which follows the gradual increase of the sequence similarity thresholds employed for the identification of these datasets. As mentioned in the “Introduction”, for reasons of completeness, size distribution of the inter-CNE distances for the four CNE collections recently identified have also been computed. The results are tabulated in the Spreadsheet within the Supplementary Data File, and the corresponding plots are given in the Supplementary Data File. They present a power-law-like pattern in their inter-CNE spacers size distribution (also found in many other CNE collections)^7^. The extent of the log-log linearity gradually increases from less conserved to ultra conserved elements, as is the case for their fractality metrics. In order to interpret these converging results, we ask for the association, if any, between the sequence similarity thresholds used in the identification of a dataset and the antiquity of the elements within this dataset. In order to address this problem, we computed the overlap of each of datasets ***(vi) – (x)*** with datasets ***(iii)*** and ***(iv)***, the latter ones comprising the ‘ancient’ sets of elements. The results are presented in the Supplementary Data File and show that the more conserved class of elements (UCNEs) overlap at a large extent with ancient elements, both Fugu and shark. Moreover, there is a gradual shift in the fraction of overlap with ancient elements as we go from the least (CNEs of similarity 75-80%) to the most (UCNEs or CNEs of similarity >95%) conserved classes of elements. This finding is compatible with the hypothesis that the proposed “Segmental-duplication / CNE elimination” model is at the origin of the observed fractality (along with self-similarity and inter-CNE size distributions following a power-law pattern) of CNE chromosomal positioning.

The underlying assumption here is that the observed gradual overlap of a series CNE collections with another dataset of known antiquity implies that the extent of this overlap expresses the order of antiquity for most of the elements belonging to the considered series of CNE collections. This assumption cannot be considered to hold true for all elements belonging to these CNE classes, as some of these elements might also be of ancient origin, though not detectable as they might have been deviated beyond recognition from ancient CNEs.

The ‘Homo – Fugu’ CNE dataset presents more developed fractality than any other CNE collection. Also, it presents the highest value for the ratio R. Notice that ‘Homo – Fugu’ dataset exceeds the fractality developed by the ‘Homo – Elephant Shark’ dataset, despite that our common ancestor with teleosts (like Fugu) is more recent than our common ancestor with cartilaginous fishes (including sharks). This result may be understood on the basis of the finding of Lee *et al.*^54^ who concluded that the jawed vertebrate ancestor had initially a great number of UCEs which have been diverged beyond recognition (i.e. they ceased to be under purifying selection) in teleosts, but survived in tetrapods. This complies with our finding that the ‘Homo – Fugu’ dataset exhibits the globally best scores as deduced by inspection of Table 1, if we take into account the predicted dependence of the extent of fractality on the intensity of the CNE loss based on the “Segmental-duplication / CNE elimination” model properties, see Figure 6 and Table 3. The finding of Lee *et al.* corresponds to type ***(iii)*** events of the model (i.e. high number of non-duplicated CNEs lost). The significance of whole genome and segmental duplications for this model also lies in accordance with the view of Wang *et al.*^55^, which correlates the numerous eliminations of UCEs that occurred in the teleost fish with the whole genome duplication that occurred in the ray-fish lineage. The role of whole genome duplication is twofold: **(a)** It makes possible the subsequent elimination of duplicated CNEs due to redundancy (type ***(ii)*** events of the model), i.e. high number of duplicated CNEs lost, expressed by a high fr value; **(b)** Simultaneously, it provides an important sequence extension which contributes to the formation of lengthy inter-CNE distances which favours fractality established at successive length scales. Convergent to the maximal fractality of ‘Homo – Fugu’ dataset herein is our previous observation studying the extent of power-law-like linearity in the distributions of inter-CNE distances where the maximum linearity is again observed in Human – Teleosts datasets^7^.

## Competing financial interests

The authors declare no competing financial interests.

## Acknowledgements

The authors would like to acknowledge Dr Philipp Bucher for fruitful discussions concerning CNE dynamics and evolution, as well as Konstantinos Apostolou – Karampelis for valuable assistance in genome analysis tasks.

## Author Contributions Statement

Y.A. and L.A. implemented the algorithms for analyzing genomic sequences using box-counting and scaling of Shannon entropy respectively. Computational analyses of CNEs using box-counting and block entropy were performed by D.P. and L.A. respectively. D.P. performed all genome analyses tasks using BEDTools and custom scripts. All authors prepared figures, contributed to the writing, as well as reviewed and approved the final version of the manuscript.

## Additional Information

-Supplementary Data File

